# Quorum Sensing Positively Regulates CPS-dependent *Autographiviridae* Phages Infection in *Vibrio alginolyticus*

**DOI:** 10.1101/2023.12.13.571448

**Authors:** Xixi Li, Chen Zhang, Shenao Li, Sixuan Liang, Xuefei Xu, Zhe Zhao

## Abstract

Quorum sensing (QS) orchestrates many bacterial behaviors, such as virulence and biofilm formation, across bacterial populations. Nevertheless, the underlying mechanism of QS regulating CPS-dependent phage-bacterium interactions remains unclear. In the present study, we report that QS upregulated the expression of CPS-dependent phage receptors, thus increasing phage adsorption and infection rates in *V. alginolyticus*. We found that QS upregulated the expression of the *ugd* gene, leading to increased synthesis of *Autographiviridae* phage receptor capsular polysaccharide (CPS) synthesis in *V. alginolyticus*. The signal molecule autoinducer-2 (AI-2) released by *V. alginolyticus* from different sources can potentially enhance CPS-dependent phage infection. Therefore, our data suggest that inhibiting quorum sensing may reduce rather than improve the therapeutic efficacy of CPS-specific phages.

**IMPORTANCE:** Phage resistance is a direct threat to phage therapy, and understanding phage-host interactions, especially bacteria block phage infection, is essential for developing successful phage therapy. In the present study, we demonstrate for the first time that *V. alginolyticus* uses QS to promote CPS-specific phage infection by upregulating the expression of *ugd*, which is necessary for the synthesis of *Autographiviridae* phage receptor capsular polysaccharide (CPS). Although increased CPS-specific phage susceptibility is a novel trade-off mediated by QS, it results in the upregulation of virulence factors, promoting biofilm development and enhanced capsular polysaccharide production in *V. alginolyticus*. This suggests that inhibiting QS may improve the effectiveness of antibiotic treatment, but it may also reduce the efficacy of phage therapy.

---

The increasing prevalence of antimicrobial resistance has led to the resurgence of phage therapy, in which bacteriophages (phages) are used to treat bacterial infections (1–3). However, because of the coevolution of phages and their host bacteria, pathogenic bacteria often resist phages, and strategies to address resistance are critical (4). However, the frequency and impact of resistance development in different bacteria vary widely (5). These phage-resistant mutant strains most typically resist phage infection by modifying phage-associated receptors on the bacterial surface that affect phage attachment (6). For example, whole-genome sequencing was performed on four spontaneous phage-resistant mutants derived from *V. cholerae* VCA0171 and found the polyQ (phage receptor) deletion resulted in phage VP1 resistance by Fan *et al*.(7). Similarly, when co-incubating strains AB900 and A9844 with their respective phages ΦFG02 and ΦCO01, the picked phage-resistant mutants harbored loss-of-function mutations in genes responsible for capsule biosynthesis, resulting in capsule loss and disruption of phage adsorption (8). In the case of *V. alginolyticus* E110, our previous data showed that five spontaneous phage-resistant mutants disrupted the adsorption of phage HH109 due to mutations in the gene synthesizing the receptor. Phage adsorption, as the first stage of bacteriophage infection, is a primary limiting factor for phage propagation and host range determination (9). Thus, focusing on the phage infection cycle’s first step is necessary to utilize the phages in bacterial disease control effectively.

Quorum sensing (QS), a cell-cell communication system, regulates gene expression based on population density and relies on the production, release, and group-wide detection of extracellular signaling molecules called autoinducers (AIs)(10, 11). *V. alginolyticus* is a prominent pathogen in the marine environment and can also act as an opportunistic pathogen for various marine animals. The strain shares a QS architecture with *V. harveyi*, which is composed of three AIs (AI-2, HAI-1, and CAI-1), their synthases, and cognate membrane-bound sensory histidine kinases (LuxM/N, LuxS/PQ, and cqsA/cqsS, respectively) (12, 13). The sRNAs, called Qrr1-5 (Quorum Regulatory RNA), together with the chaperone Hfq, regulate the translation of the master QS regulator (MQSRs) AphA and LuxR (14). An additional QS-mediated system comprises the small molecule nitric oxide (NO) and heme-nitric oxide/oxygen (H-NOX) associated histidine kinase HahK. This system allows *V. harveyi* to sense NO and regulate gene expression through a QS pathway that includes AphA and LuxR (15). The expression of *aphA* in the QS circuits of vibrios is negatively associated with *luxR* levels. AphA is specifically responsible for regulating gene expression during the low-cell-density (LCD) stage, while LuxR is involved in the high-cell-density (HCD) stage (16, 17). In *V. harveyi*, the combined or individual actions of these two MQSRs control the expression of approximately 600 genes, leading to the emergence of different cooperative behaviors such as bioluminescence, biofilm formation, motility, and virulence (18).

It was recently demonstrated that QS is closely related to phage infection and could influence bacterial antiphage defenses by reducing the number of phage receptors on the cell surface. *Pseudomonas aeruginosa* modulates its sensitivity to phage vB_Pae_QDWS infection via a mechanism that QS upregulates the expression of *galU* to increase the yield of the phage receptor lipopolysaccharide (LPS) (19). However, *V. cholerae* phage JSF35 receptor LPS O-antigen was downregulated when supplemented with the autoinducers CAI-1 or AI-2 (20). Besides, *V. anguillarum* QS used N-acyl homoserine lactones (AHLs) as signal molecules to downregulate KVP40 phage receptor *OmpK* expression (21). These research advances suggest that host bacterial QS signaling regulates the expression of many phage receptors with different regulatory outcomes. In addition, QS regulates the expression of the CRISPR-Cas immune system and activates cas gene expression in *P. aeruginosa* and *Serratia*, which protects the bacteria against phage infection (22, 23). Therefore, Manipulation of QS may impact the level of host resistance and the expression of various phage receptors during infection. For example, these QS systems activate the CRISPR-cas immune system’s expression and type IV pili, the receptor for many Pseudomonas phages (22, 24). It is unknown whether QS plays a role in CPS-dependent phage infection by modulating phage adsorption or host resistance in *V. alginolyticus*.

Our previous work identified CPS as the receptors for *Autographiviridae* phages HH109. Here, we characterized two other *Autographiviridae* phages, φVP505 and φVP506, which infect their host in a CPS-dependent manner. Our data demonstrated that QS upregulates the expression of *ugd*, a key gene for CPS synthesis, leading to an increased rate of *V. alginolyticus* phage adsorption and, thus, increased phage infection.

Overall, these results are highly relevant in the context of phage therapy, as QS inhibition may reduce the therapeutic effect of the phage system. The discovery may increase the efficiency of phage control of vibriosis.

## RESULTS

### Quorum sensing influences phage HH109 infection efficiency

We investigated the impact of the QS system on phage HH109 resistance. Phage HH109 significantly reduced the cell density in the cultures of wild-type E110 and QS-deficient mutants Δ*luxS*, Δ*luxM*, Δ*cqsA*, and Δ*hnoX* compared to that in control cultures without the phage. However, Δ*luxS*, Δ*luxM*, Δ*cqsA*, and Δ*hnoX* exhibited a higher cell density within 4 h during the incubation period than E110 (Fig. 1A-D). Meanwhile, to explore the effects of signaling molecules on *V. alginolyticus* QS-deficient mutants encountering phage HH109 in communities, we administered cell-free culture fluids from *V. alginolyticus* E110, which produce all QS signaling molecules. Remarkably, cell-free culture fluids from *V. alginolyticus* E110 completely restored cell lysis of mutants Δ*luxS*, Δ*luxM*, Δ*cqsA* by the phage HH109 (Fig. 1A-C). By contrast, cell-free culture fluids from E110 did not affect cell lysis of mutants Δ*hnoX* HH109 relative to the medium alone (Fig. 1D). The wild-type E110 and the complemented strain Δ*luxS:* pBBR1-*luxS*, Δ*luxM:* pBBR1-*luxM*, Δ*cqsA:* pBBR1-*cqsA*, Δ*hnoX:* pBBR1-*hnoX* exhibited similar cell density (Fig. S1A-D). These results suggest that the QS positively regulates phage HH109 sensitivity of *V. alginolyticus* E110.

**Figure 1.**
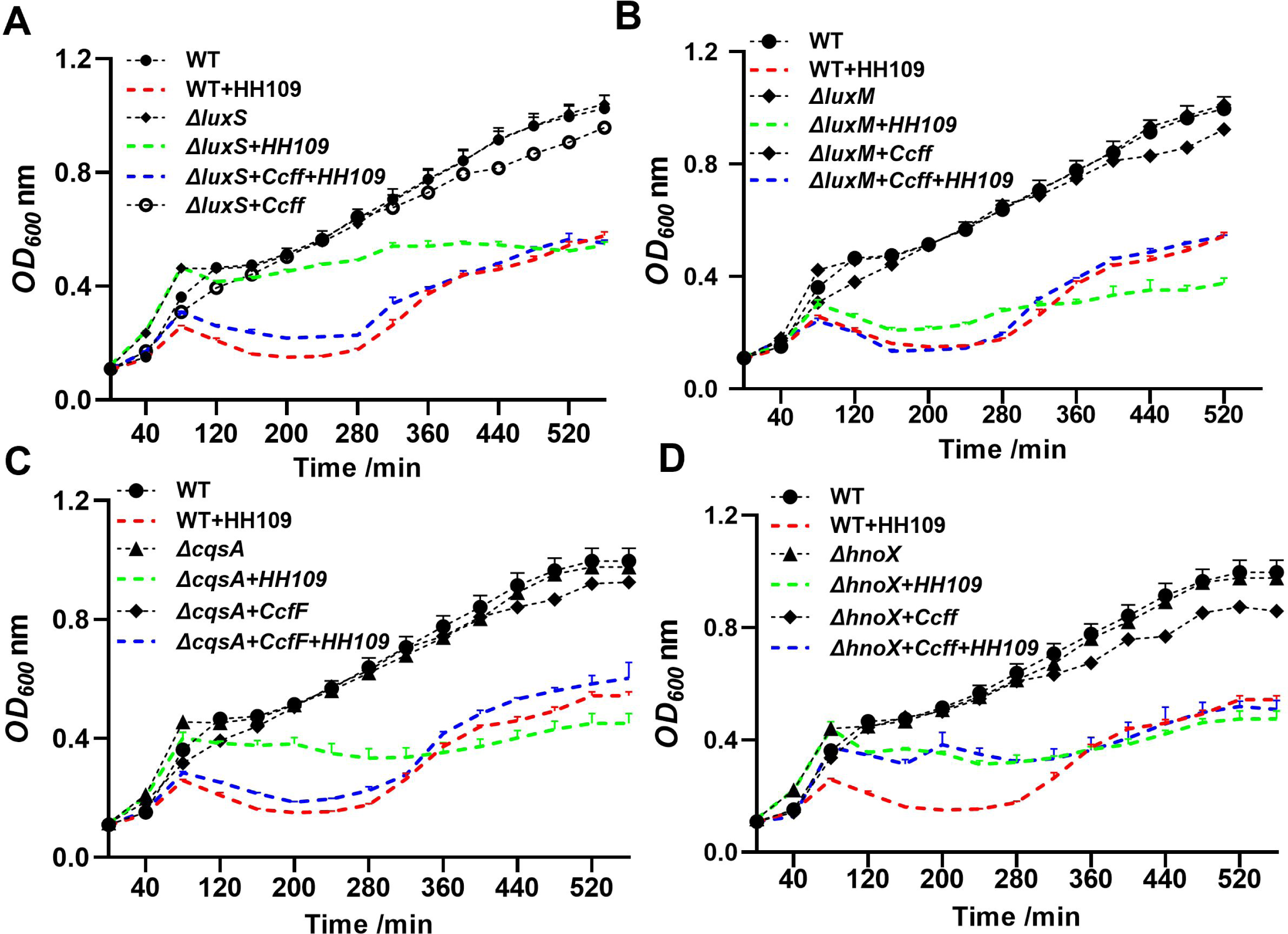
*V. alginolyticus* E110 quorum-sensing (QS) mutants influence phage infection capacity. **(A)** Optical densities (OD_600_) of QS mutants Δ*luxS* grown in a medium containing 2% NaCl and in cell-free culture fluids (Ccff) from E110 wild-type (WT) in the presence or absence of phage HH109 at a multiplicity of infection (MOI) of 0.001 were measured in a 96-well microtiter plate containing 200 μL of each culture using the Bioscreen C at every 40-min interval for 520 min. WT E110 was grown in an LB medium containing 2% NaCl as a control. Data are averages of three samples with standard deviations (error bars). **(B)** OD_600_ of QS mutants Δ*luxM* grown in a medium containing 2% NaCl and in cell-free culture fluids (Ccff) from E110 (WT) in the presence or absence of phage HH109 at an MOI of 0.001 were measured in a 96-well microtiter plate containing 200 μL of each culture using the Bioscreen C at every 40-min for 520 min. E110 (WT) was grown in an LB medium containing 2% NaCl as a control. Data are averages of three samples with standard deviations (error bars). **(C)** OD_600_ of QS mutants Δ*cqsA* grown in a medium containing 2% NaCl and in cell-free culture fluids (Ccff) from E110 WT in the presence or absence of phage HH109 at an MOI of 0.001 were measured in a 96-well microtiter plate containing 200 μL of each culture using the Bioscreen C at every 40-min interval for 520 min. WT E110 was grown in an LB medium containing 2% NaCl as a control. Data are averages of three samples with standard deviations (error bars). **(D)** OD_600_ of QS mutants Δ*hnoX* grown in a medium containing 2% NaCl and in cell-free culture fluids (Ccff) from E110 WT in the presence or absence of phage HH109 at an MOI of 0.001 were measured in a 96-well microtiter plate containing 200 μL of each culture using the Bioscreen C at every 40-min for 520 min. WT E110 was grown in an LB medium containing 2% NaCl as a control. Data are averages of three samples with standard deviations (error bars).

### Quorum sensing influences phage HH109 adsorption

Next, to investigate the mechanisms associated with the altered susceptibility of *V. alginolyticus* E110 strains to phage infections, we measured whether QS-deficient mutants affect phage adsorption. The QS-deficient mutants Δ*luxS*, Δ*luxM*, Δ*cqsA*, and Δ*hnoX* exhibited a significant reduction in phage adsorption rates compared to that of the E110 wild-type strain. However, the adsorption rates were higher for the complemented strains and QS-deficient strains Δ*luxS*, Δ*luxM*, and Δ*cqsA* grown in cell-free culture fluids from E110 than that of E110 wild-type strain (Fig. 2A-C). Furthermore, the cell-free medium from E110 did not enhance the adsorption of HH109 to mutant Δ*hnoX* (Fig. 2D). This revealed that QS positively regulated phage susceptibility by increasing the phage adsorption rate.

**Figure 2.**
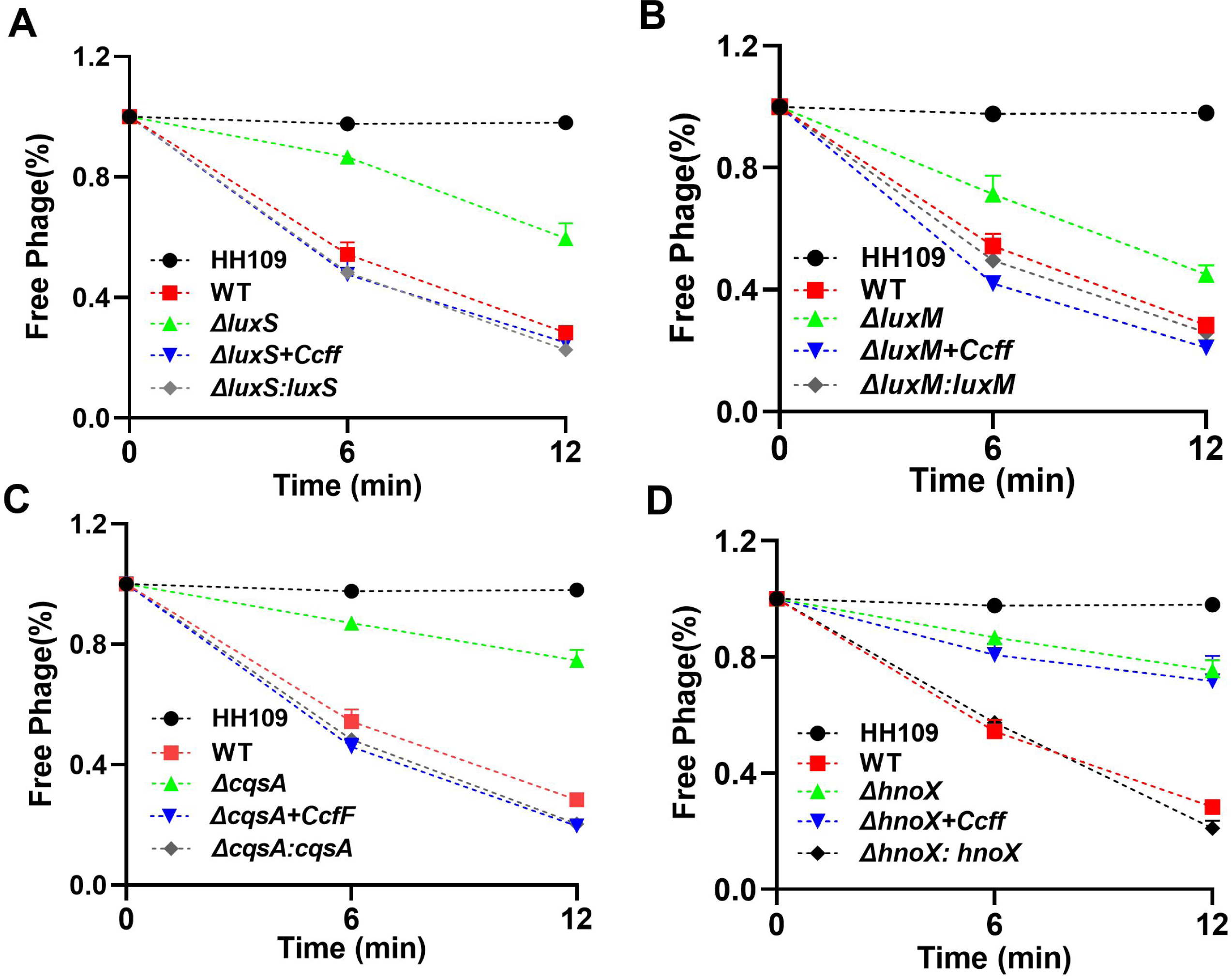
Adsorption rate of phages by its host strains *V. alginolyticus* E110 and quorum-sensing (QS) mutants. **(A)** Adsorption rate of phage HH109 by its host strains *V. alginolyticus* E110 and QS mutants Δ*luxS and* complemented strain Δ*luxS: luxS* at every 6-min interval for 12 min. Data are averages of three samples with standard deviations (error bars). **(B)** Adsorption rate of phage HH109 by its host strains *V. alginolyticus* E110 and QS mutants Δ*luxM* and complemented strain Δ*luxM: luxM* at every 6-min interval for 12 min. Data are averages of three samples with standard deviations (error bars). **(C)** Adsorption rate of phage HH109 by its host strains *V. alginolyticus* E110 and QS mutants Δ*cqsA* and complemented strain Δ*cqsA*: *cqsA* at every 6-min interval for 12 min. Data are averages of three samples with standard deviations (error bars). **(D)** Adsorption rate of phage HH109 by its host strains E110 and QS mutants Δ*hnoX* and complemented strain Δ*hnoX: hnoX* at every 6-min interval for 12 min. Data are averages of three samples with standard deviations (error bars).

### Quorum sensing influences *Autographiviridae* phage receptor expression

QS can regulate numerous bacterial behaviors, such as bioluminescence, motility, biofilm formation, virulence factor expression, or exopolysaccharide (EPS) production (25–28). Hence, we envisaged phage receptors are regulated by QS, which would cause them to become more resistant to phage infection. We first constructed aphA and luxR knockout mutant strains of *V. alginolyticus* E110 to test the hypothesis. The QS regulator AphA and LuxR is controlled and regulated by *luxM*, *luxS*, and *cqsA* produced autoinducers HAI-1, AI-2, and CAI-1, respectively, sensed by corresponding histidine kinases (29, 30). This analysis showed that compared to the wild-type control group, *V. alginolyticus* mutant Δ*luxR* exhibited increased phage infection resistance and prolonged the adsorption time of phage HH109 (Fig. 3A-B). However, the mutant Δ*aphA* showed little change (Fig. S2A-B). Those results further confirm that QS promotes phage HH109 infection assay. Secondly, we quantified the yield of CPS for mutants Δ*aphA* and Δ*luxR* using a standard curve of pure standard glucuronolactone (Sigma-Aldrich) (Fig. 3C). The results showed a decrease in CPS production in *V. alginolyticus* Δ*luxR* QS mutant compared to the wild type strains. Conversely, the Δ*aphA* QS mutant exhibited CPS production similar to the control strains. Furthermore, the complementing strain Δ*luxR*: *luxR* displayed CPS production nearly identical to the wild-type strains (Fig. 3D). These results suggest that QS can regulate the synthesis of receptor CPS to combat *Autographiviridae* phage infection.

**Figure 3.**
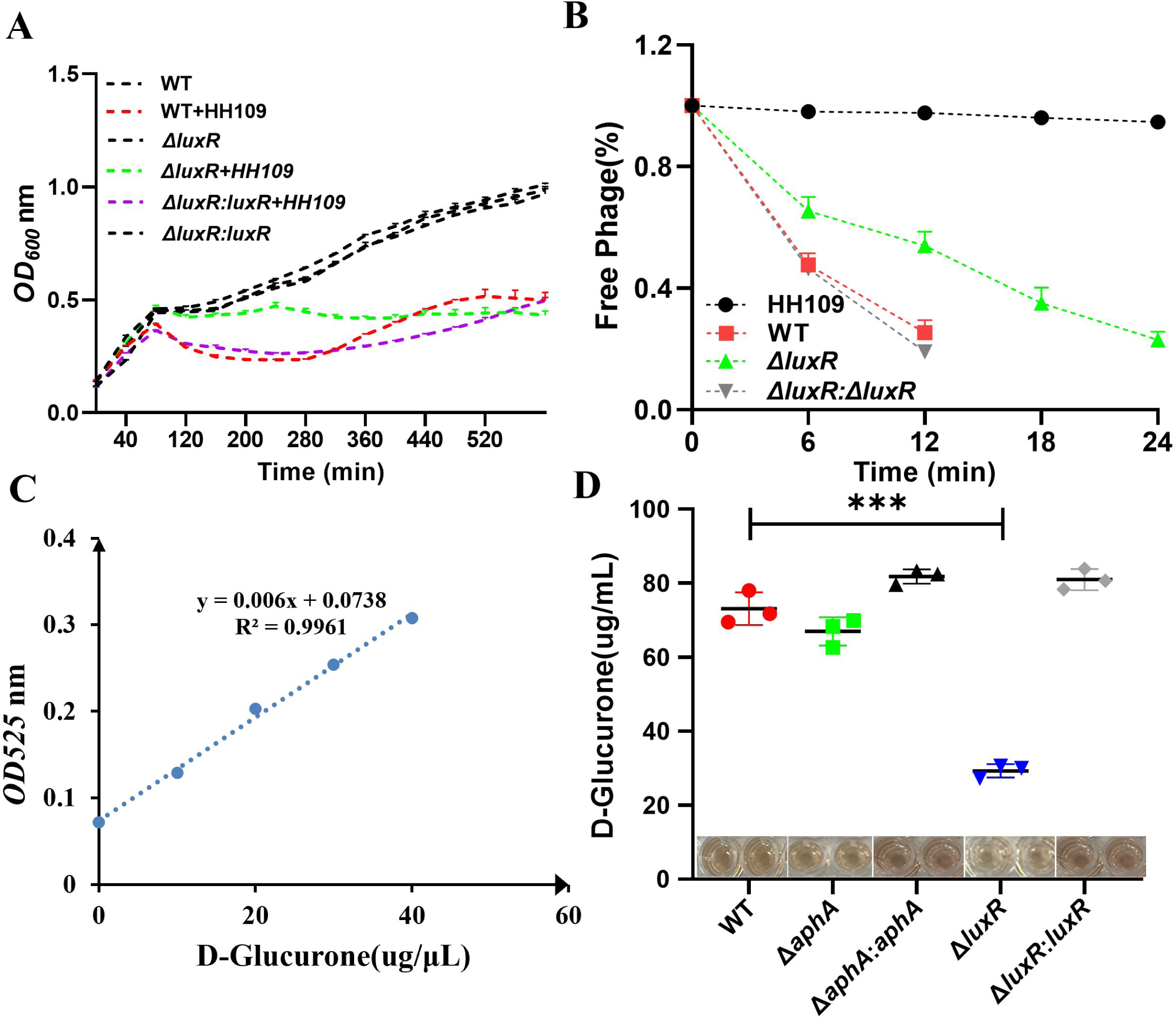
Quorum-sensing Regulator luxR Directly influences *Autographiviridae* phage HH109 infection capacity Through Reduced capsule production in *V. alginolyticus* E110. **(A)** Optical densities (OD_600_) of cultures of E110 (WT) and mutants Δ*luxR* and complemented strain Δ*luxR:*Δ*luxR* in the presence or absence of phage HH109 at a multiplicity of infection (MOI) of 0.001 were measured in a 96-well microtiter plate containing 200 μL of each culture using the Bioscreen C at every 40-min for 520 min. Data are averages of three samples with standard deviations (error bars). **(B)** Adsorption rate of phage HH109 by its host strains E110 (WT) and mutants Δ*luxR* and complemented strain Δ*luxR:*Δ*luxR* at every 6-min interval for 24 min. Data are averages of three samples with standard deviations (error bars). **(C)** Standard curve for the Uronic acid lactone capsule quantification assay, calibrated with glucuronolactone standards ranging from 0 to 40 μg/mL. OD_525_, optical density at 525 nm. **(D)** Production of CPS for *V. alginolyticus* E110 and QS mutants Δ*aphA* or Δ*luxR* and complemented strain Δ*aphA:aphA* or Δ*luxR: luxR* at OD_600_ 1.5. Data are averages of three samples with standard deviations (error bars). ***, *P* < 0.001 (paired *t* test).

To further verify the above conclusion, we characterized two other *Autographiviridae* phages, φVP505 and φVP506, which infect the host *V. alginolyticus* VP505 and VP506, respectively, in a CPS-dependent manner. As shown in Fig. 4, deletions of *ugd* and *wecA*, two key genes for CPS synthesis, caused an increase in host bacterial resistance (Fig. 4A) and a significant reduction in the phage adsorption (Fig. 4B) compared with wild-type strains. Subsequently, VP505Δ*aphA*, VP505Δ*luxR*, VP506Δ*aphA*, and VP506Δ*luxR* deletion mutants of *V. alginolyticus* VP505 and VP506 were constructed, respectively. Phage infection and adsorption experiments showed that compared to the wild-type and complemented strains VP505Δ*luxR*: *luxR* and VP506Δ*luxR*: *luxR*, CPS-dependent phages φVP505 and φVP506 infect their QS mutants VP505Δ*luxR* and VP506Δ*luxR* more rapidly produced phage-resistant strains and had longer adsorption times, respectively (Fig. 5A-D). However, the VP505Δ*aphA* and VP506Δ*aphA* mutants showed almost no changes (Fig. S2C-F). Furthermore, capsule quantification results showed that CPS production was reduced in VP505Δ*luxR* and VP506Δ*luxR* QS mutants. Conversely, the CPS production of the VP505Δ*aphA* and VP506Δ*aphA*-QS mutants was similar to that of the control strain and the complemented strains VP505Δ*luxR*: *luxR* and VP506Δ*luxR*: *luxR*, respectively (Fig. 5E-F). These data, therefore, collectively confirm that QS-controlled CPS-dependent group behavior in *V. alginolyticus*.

**Figure 4.**
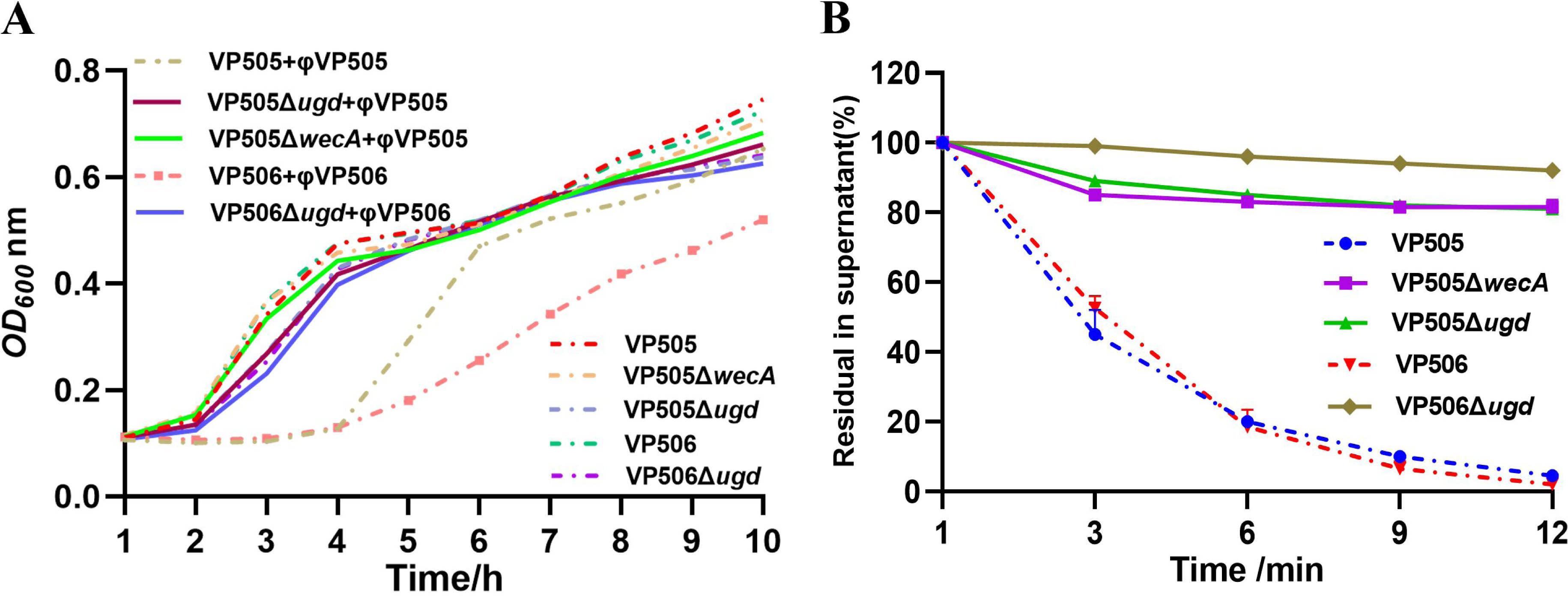
Identification of CPS as an essential receptor for *V. alginolyticus* phage φVP505 and φVP506 infection. **(A)** Bacterial growth was monitored by measuring the turbidity (OD_600_) of wild-type and two deletion mutants with or without the phage φVP505 and φVP506. Data are displayed as the means ± SD from three independent experiments. **(B)** Phage adsorption assays. Another phage, φVP505, and φVP506, were introduced to verify that the CPS of *V. alginolyticus* serves as a receptor for infection of the phages. Two deletion mutants, Δ*wecA* and Δ*ugd*, were generated in *V. alginolyticus* strain VP505, and one deletion mutant Δ*ugd* in *V. alginolyticus* strain VP506 was subjected to adsorption assay. The samples were taken at the indicated times, and residual phages were in the supernatant. Data are displayed as the means ± SD from three independent experiments.

**Figure 5.**
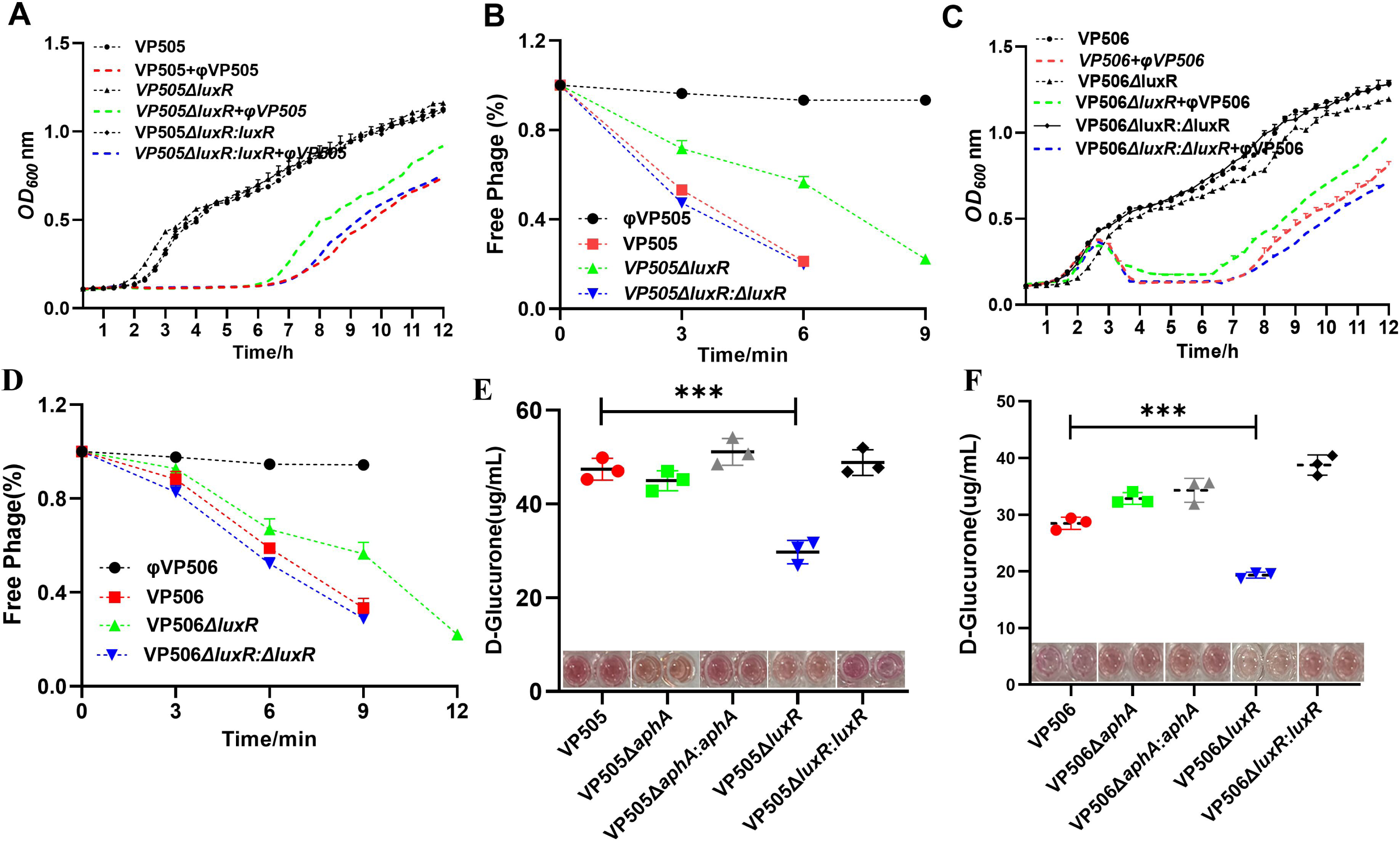
Quorum-Sensing Regulator luxR Directly influences *Autographiviridae* phages φVP505 and φVP506 infection capacity and capsule production in *V. alginolyticus* VP505 and VP506. **(A)** OD_600_ of cultures of VP505 and QS mutants VP505Δ*luxR* and complemented strain VP505Δ*luxR:*Δ*luxR* in the presence or absence of phage φVP505 at a multiplicity of infection (MOI) of 0.01 were measured in a 96-well microtiter plate containing 200 μL of each culture using the Bioscreen C at every 20-min interval for 12 h. Data are averages of three samples with standard deviations (error bars). **(B)** Adsorption rate of phage φVP505 by its host strains *V. alginolyticus* VP505 and QS mutants VP505Δ*luxR* and complemented strain VP505Δ*luxR:*Δ*luxR* at every 3-min for 9 min. Data are averages of three samples with standard deviations (error bars). **(C)** OD_600_ of cultures of VP506 and QS mutants VP506Δ*luxR* and complemented strain VP506Δ*luxR:*Δ*luxR* in the presence or absence of phage φVP506 at a multiplicity of infection (MOI) of 0.001 were measured in a 96-well microtiter plate containing 200 μL of each culture using the Bioscreen C at every 20-min interval for 12 h. Data are averages of three samples with standard deviations (error bars). **(D)** Adsorption rate of phage φVP506 by its host strains *V. alginolyticus* VP506 and QS mutants VP506Δ*luxR* and complemented strain VP506Δ*luxR:*Δ*luxR* at every 3-min interval for 12 min. Data are averages of three samples with standard deviations (error bars). **(E)** Production of CPS for *V. alginolyticus* VP505 and QS mutants VP505Δ*aphA* and complemented strain VP505Δ*aphA: aphA* at OD_600_ 1.5. Data are averages of three samples with standard deviations (error bars). ***, *P* < 0.001(paired *t* test). **(F)** Production of CPS for *V. alginolyticus* VP506 and QS mutants VP506Δ*aphA* and complemented strain VP506Δ*aphA: aphA* at OD_600_ 1.5. Data are averages of three samples with standard deviations (error bars). ***, *P* < 0.001 (paired *t* test).

### *Ugd* expression is activated by Quorum sensing

Our previous study revealed that the CPS gene cluster of *V. alginolyticus* E110 comprises 36 related genes, of which only five genes (*gtr1*, *wecA*, *orf2*, *gtr6*, and *ugd*) were related to phage HH109 infection. To this end, we investigated *gtr1*, *wecA*, *orf2*, *gtr6*, and *ugd* expression in *V. alginolyticus* strains. When the regulator aphA was deleted, resulting in cells corresponding to a high-density phase, the expression levels of *gtr1* and *gtr6* were significantly increased. However, the expression of *wecA*, *orf2*, and *ugd* did not change compared to the wild-type strain E110. On the other hand, when the regulator luxR was deleted, resulting in cells corresponding to a low-density phase, only the expression level of *ugd* was significantly decreased compared to the wild-type strain E110 (Fig. 6A). It is important to note that the expression of *ugd* is dependent on the strain growth phase, with higher expression at high cell densities compared to low cell densities (Fig. 6B and Fig. S3B). Similar results were further confirmed in *V. alginolyticus* VP505 and VP506. Compared with the wild-type strain, the expression level of *ugd* in the *V. alginolyticus ΔluxR*-QS mutants (VP505Δ*luxR* and VP506Δ*luxR*) was significantly reduced. In contrast, the expression level of *ugd* in the *V. alginolyticus ΔaphA*-QS mutants (VP505Δ*aphA* and VP506Δ*aphA*) was not changed (Fig. S3A). Therefore, we can conclude that the expression of *ugd* is regulated by QS in *V. alginolyticus*.

**Figure 6.**
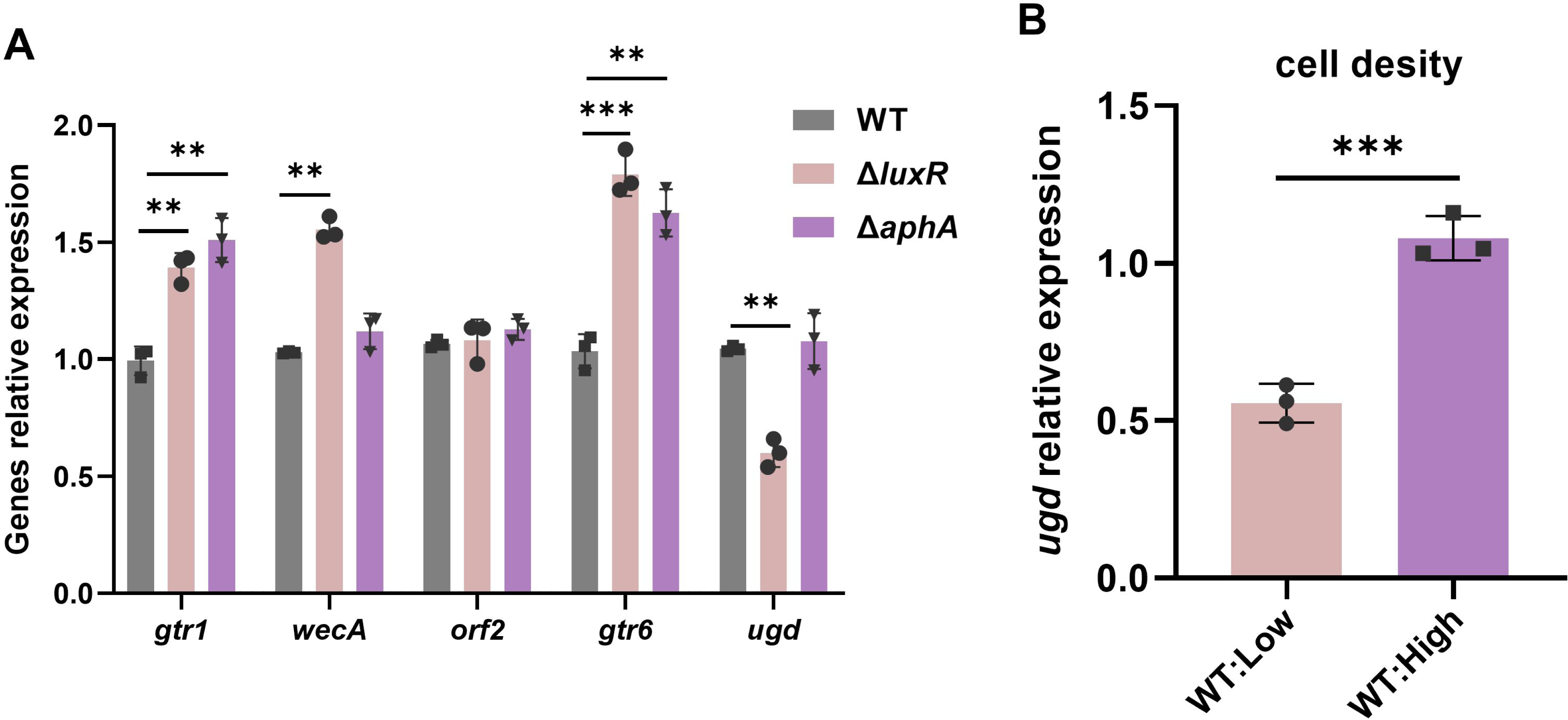
QS activates *ugd* expression. **(A)** The expression of key genes of CPS synthesis at high cell density in wild-type (WT) E110 and the designated QS mutants. **(B)** Relative *ugd* expression was measured by RT-qPCR in *V. alginolyticus* E110 cells at low and high cell densities (OD_600_, 0.8, and 2.5, respectively). The reference gene was *dnaK*. Data are averages of three samples with standard deviations (error bars). **, *P* < 0.01; ***, *P* < 0.001(paired *t* test).

### Signaling molecule autoinducer-2 released by multiple *V. alginolyticus* hastens phage HH109 infection

The signal molecule autoinducer-2 (AI-2) synthesized by the LuxS has been recognized as a universal language for cell-to-cell communication (31). To test the effect of AI-2 released by *V. alginolyticus* from different sources on the infection process of phage HH109, we administered cell-free culture fluids harvested from double-mutants Δ*luxM*Δ*cqsA V. alginolyticus* E110, VP505, and VP506, which higher cell density produced more AI-2 activity within 24 h (Fig. S4). Identical to the case of double-mutants Δ*luxM*Δ*cqsA V. alginolyticus* E110 cell-free culture fluids, the addition of cell-free culture fluids from double-mutants Δ*luxM*Δ*cqsA V. alginolyticus* VP505 and VP506 promote the adsorption and lysis of phage HH109 on host *V. alginolyticus* E110. By contrast, the phage HH109 did not reach the basal level of adsorption and lysis for mutants Δ*luxS V. alginolyticus* E110 cultured in a conventional culture medium, compared to the wild-type control (Fig. 7A-C). These findings suggest that AI-2, released by *V. alginolyticus* from different sources, can potentially enhance CPS-dependent phage HH109 infection.

**Figure 7.**
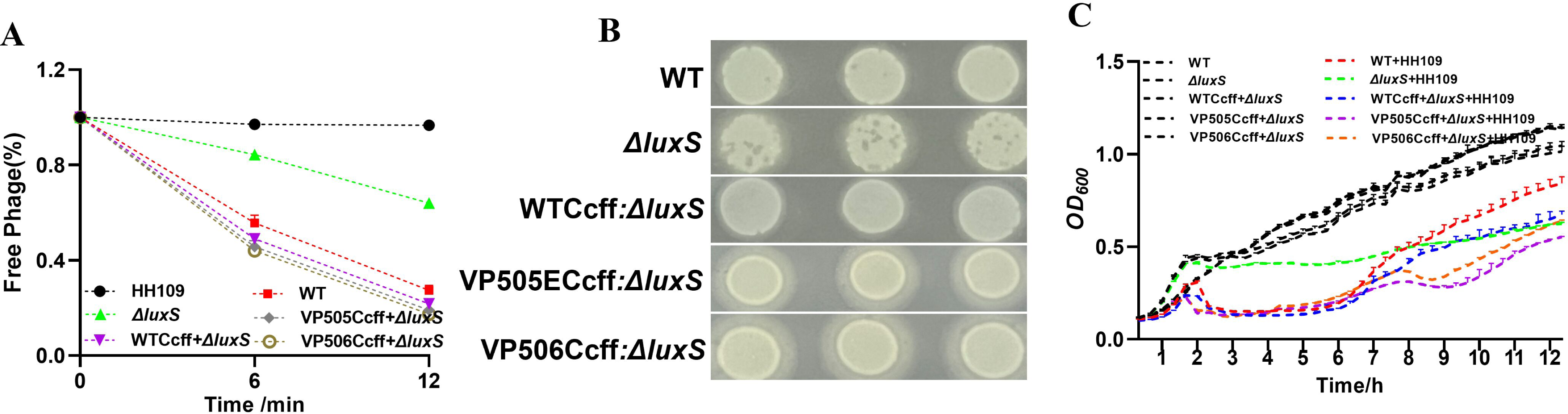
*V. alginolyticus* cell-free culture fluids inhibit the fitness of quorum-sensing (QS) mutants Δ*luxS* in the presence of phage HH109. **(A)** Adsorption rate of phage HH109 by its host strains *V. alginolyticus* E110 (WT) and Δ*luxS V. alginolyticus* E110 grow in LB with 2% NaCl or LB with 2% NaCl containing cell-free culture fluids from *V. alginolyticus* E110 (WTCcff), or cell-free culture fluids from *V. alginolyticus* VP505 (VP505Ccff), or cell-free culture fluids from *V. alginolyticus* VP506 (VP506Ccff). Data are averages of three samples with standard deviations (error bars). **(B)** Cleavage efficiency of supernatant from phage HH109 adsorbed E110 (WT) and Δ*luxS V. alginolyticus* E110 for 12 min on host E110 at 5 h by spotting assays. E110: The infectivity of the supernatant (non-adsorbed phage) of 3.6×10^6^ Pfu/mL phage HH109 adsorbed on host E110 (growth in LB with 2% NaCl for 24 h) for 12 min. Δ*luxS*: The infectivity of the supernatant (non-adsorbed phage) of 3.6×10^6^ Pfu/mL phage HH109 adsorbed on host E110 (growth in LB with 2% NaCl for 24 h) for 12 min. WTCcff*+*Δ*luxS*: The infectivity of the supernatant (non-adsorbed phage) of 3.6×10^6^ Pfu/mL phage HH109 adsorbed on host E110 (growth in LB with 2% NaCl containing cell-free culture fluids from *V. alginolyticus* E110 for 24 h) for 12 min. VP505Ccff*+*Δ*luxS*: The infectivity of the supernatant (non-adsorbed phage) of 3.6×10^6^ Pfu/mL phage HH109 adsorbed on host E110 (growth in LB with 2% NaCl containing cell-free culture fluids from *V. alginolyticus* VP505 for 24 h). VP506Ccff*+*Δ*luxS*: The infectivity of the supernatant (non-adsorbed phage) of 3.6×10^6^ Pfu/mL phage HH109 adsorbed on host E110 (growth in LB with 2% NaCl containing cell-free culture fluids from *V. alginolyticus* VP506 for 24 h). **(C)** Optical densities (OD_600_) of cultures of E110 wild-type (WT) and Δ*luxS V. alginolyticus* E110 grow in LB with 2% NaCl or LB with 2% NaCl containing cell-free culture fluids from *V. alginolyticus* E110 (E110Ccff), or cell-free culture fluids from *V. alginolyticus* VP505 (VP505Ccff), or cell-free culture fluids from *V. alginolyticus* VP506 (VP506Ccff). Data are averages of three samples with standard deviations (error bars).

## DISCUSSION

Considering the recent discovery that bacteria can utilize QS to reduce phage receptor expression under conditions of high infection risk (32), we aimed to investigate the role of QS in modulating the sensitivity of *V. alginolyticus* strains to CPS-dependent phage. The reduced susceptibility to *Autographiviridae* phages in *V. alginolyticus* QS-deficient mutants provided the first indications of a QS-regulated mechanism of phage protection. Our subsequent studies of reduced CPS production and low expression gene *ugd*, a key CPS synthesis gene in *V. alginolyticus* QS-deficient mutants. A schematic representation of the proposed mechanism is shown in Figure 8. Our results suggest that *V. alginolyticus* QS regulates *ugd* expression, which is required for CPS receptor synthesis, and subsequently affects the susceptibility of *V. alginolyticus* to HH109, φVP506, and φVP506 phages infection.

**Figure 8.**
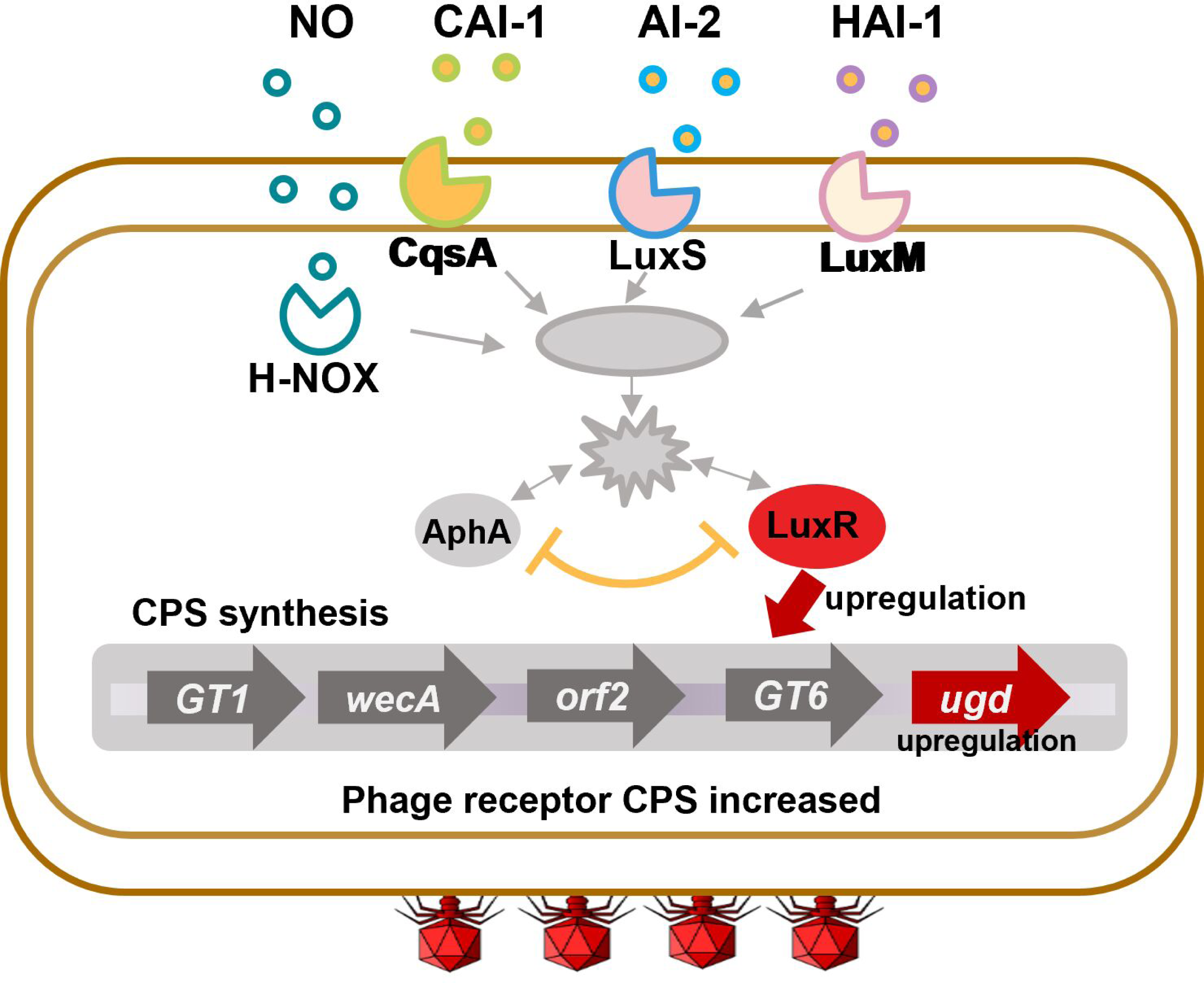
Schematic representation of the mechanism by which las QS regulates the resistance of *V. alginolyticus* E110 to phage HH109. The QS positively regulates the expression of *ugd*, which is involved in CPS biosynthesis, thereby promoting phage adsorption.

Quorum sensing upregulated the expression of many critical features of opportunistic pathogens, including virulence gene expression and the CRISPR-Cas immune system that protects the bacteria from phage infection (22, 23). In contrast, our results suggest that QS positively regulates *Autographiviridae* phage susceptibility in *V. alginolyticus* cells and may improve the efficacy of phage therapy. In agreement with our results, previous reports found that inhibition of QS may reduce the therapeutic effect of phage vB_Pae_QDWS(19). Broniewski reported that chemical inhibition of QS reduces phage adsorption rate by downregulating type IV pili, which causes delayed lysis of bacterial cultures and promotes the development of CRISPR immunity (33). To this end, QS may have dual effects of reducing phage adsorption rate and promoting CRISPR immune development in *V. alginolyticus*. However, genome sequencing results indicated that the CRISPR-Cas system does not exist in *V. alginolyticus* E110, VP505, and VP506. QS may have different effects on phage infection in these strains. Once more, our findings revealed elevated CPS-specific phage adsorption caused by *V. alginolyticus* QS, a phenomenon distinct from previous studies on *V. anguillarum*, *V. cholerae*, and *Escherichia coli* (32, 34, 35). However, it is worth noting that numerous *Pseudomonas* phages identify type IV pili, which are known to be positively controlled by QS. This aligns with our experimental outcomes (24, 33, 36). Therefore, it is evident that the regulation of phage resistance through QS manifests remarkably diverse and intricate.

Capsule polysaccharides, as the primary receptors for *Autographiviridae* phages HH109, φVP506, and φVP506, require multiple genes to participate in their synthesis in *Vibrios* (37). However, our previous research showed that only the deletion of *gtr1*, *wecA*, *orf2*, *gtr6*, and *ugd* CPS synthesis genes specifically impacts the lysis of phage HH109 on the *V. alginolyticus* host E110. Transcriptional analysis was performed to examine the gene expression involved in phage infection. The results revealed a significant correlation between *ugd* expression and QS. However, no correlation was observed between the expression of *gtr1*, *wecA*, *orf2*, and *gtr6* and QS in *V. alginolyticus* (Fig. 6). *V. alginolyticus* LuxR, a key quorum-sensing regulator, controls the expression of ∼280 genes and regulates important virulence-associated factors, such as CPS production, motility, and biofilm formation (38, 39). Thus, the coordination is achieved through QS pathways and leads to an increase in the expression of *ugd*, a key gene for CPS synthesis, and an increased production of CPS at high bacterial cell density. The virulence factor CPS is a phage receptor and one of the factors involved in biofilm formation, representing a barrier between the phage and its receptor(8, 40). It is worth noting that the synthesis pathway of CPS is positively regulated by QS, suggesting that there may be a trade-off between phage sensitivity and CPS-based bacterial virulence. Our data indicate that inhibiting quorum sensing may reduce rather than improve the therapeutic efficacy of CPS-specific phages, which is likely a general feature when QS positively regulates phage receptors.

Bacteria typically exist in mixed microbial communities with other QS bacteria in natural environments. AI-2 acts as a crucial signal molecule for interspecific communication of Vibrios that regulate many important physiological functions of bacteria, and its synthesis and secretion are regulated by the cascade of *luxS*, *luxR*, and *aphA* genes (16). In most bacteria, the activity of AI-2 reaches its highest point during the mid-to-late logarithmic growth phase and then decreases rapidly as the stationary phase sets in (41). In contrast, strain E110 exhibits a distinct characteristic in which its AI-2 remains active even in the stable phase (Fig. S4). This observation aligns with the AI-2 signaling molecule activity found in *V. ichthyoenteri* (42). The molecular structure of AI-2 may vary among different bacteria. However, researchers widely believed that AI-2 could be a derivative of DPD (4, 5-dihydroxy-2, 3-pentanedione) (43, 44). Our findings demonstrate that externally added cell-free culture fluids from double-mutants Δ*luxM*Δ*cqsA V. alginolyticus* E110, VP505, and VP506 facilitate the adsorption and lysis of phage HH109 on host *V. alginolyticus* E110 (Fig. 7). However, using the method of directly adding cell-free culture supernatant to study the effect of AI-2 has several interference factors on the test due to the complex composition of the supernatant. Moreover, it is currently unavailable to obtain commercial signal molecule AI-2 in China, and preparing synthetic AI-2 through recombinant expression is time-consuming and challenging. This discovery suggests that synthetic molecules designed for bacterial QS systems may have unintended consequences for applying bacteriophages to treat resistant strains.

To successfully apply phage therapy in the treatment of vibriosis, it is crucial to have a comprehensive understanding of the interactions between phages and their hosts and the mechanisms regulating antiphage strategies in *Vibrio*. Our findings revealed that the regulation of CPS-specific phage susceptibility in *V. alginolyticus* is heavily influenced by QS. We observed a positive correlation between las QS and *ugd*, which is involved in the synthesis of CPS. Additionally, our results demonstrated that AI-2 derived from diverse sources of *V. alginolyticus* promotes the infection of bacteriophage HH109, indicating the potential of utilizing QS signaling molecules as a promising approach in future antimicrobial therapy.

## MATERIALS AND METHODS

### Bacterial strains, phages, plasmids, and growth conditions

The bacterial strains and plasmids used in the present study are detailed in Table S1. *Escherichia coli* strains were grown in Luria-Bertani broth (LB, Huankai, China) or Luria-Bertani agar (LB, Huankai, China) with aeration at 37°C. *V. alginolyticus* strains and derivatives were typically grown at 30°C in an LB medium supplemented with 2% NaCl. Antibiotics were used at the following concentrations: chloramphenicol at 5 μg/mL (Cm 5) for *V. alginolyticus* and 25 μg/mL (Cm 25) for *E. coli*. The 0.2% D-Glucose and 0.2% L- (+)-Arabinose purchased from Sigma were used to screen deletion mutant strains. 300 μM 2,6-diaminopimelic acid (DAP) was used to grow auxotrophic *E. coli* GEB833.

The *Autographiviridae* phage φVP505 and φVP506 were kindly provided by Professor Mao Lin (Jimei University). In this study, φVP505 was propagated on host *V. alginolyticus* VP505 with an optimal MOI of 0.01. On the other hand, φVP506 grew on *V. alginolyticus* VP506 with an optimal MOI of 0.1. The *Autographiviridae* phage HH109, an obligate lytic phage, was previously isolated and characterized in our laboratory as previously described (45).

### Gene knockout and complementation

All gene deletion strains were generated using homologous recombination, as described previously (46). The primers used for the inactivation of QS-related genes are listed in Table S2. The QS-deficient mutants were selected using colony PCR. The complemented strains of QS-deficient mutants were constructed by transforming a QS-expressing plasmid with Cm 25 to the QS-deficient mutants.

### Lysis curve determination

Transfer 100 μL of *V. alginolyticus* or QS-complemented bacteria and 10 μL of phage with an MOI of 0.01, 0.1, or 0.001 into fresh LB medium containing 2% NaCl, and then aliquot them into 96-well plates while transferring the same proportion of QS-deficient mutants and phage into the cell-free culture fluids. Growth kinetics of wild-type, deletion, and complementary strains were determined using Bioscreen C (FP-1100-C, Bioscreen) measuring OD_600_ at every 20-min interval.

*V. alginolyticus* cells were grown overnight in an LB containing 2% NaCl with shaking at 30°C. The cells were pelleted by centrifugation for 2 min at 10,000 rpm, and the resulting cell-free fluids were filtered through 0.22 μm filters (Millipore, USA). Unless otherwise stated, cell-free culture fluids were diluted 1:3 with fresh medium to culture *V. alginolyticus* QS-deficient mutants in this study.

### Phage Adsorption Assays

*V. alginolyticus* and QS-deficient mutants were inoculated in cell-free culture fluids or fresh LB medium with 2% NaCl until the OD_600_ reached 2.0. Bacterial cells and 10-fold dilution phage were mixed at an MOI of 0.01 or 0.1. The mixtures were incubated at 30°C for 3, 6, 9, 12, 18, and 24 min, and then were centrifuged at 4°C for 1 min. Subsequently, the double-layer agar plate method quantified free phage particles in the supernatant (45). To visualize the effect of AI-2 on the adsorption of the QS mutant Δ*luxS* by phage HH109, a phage drop assay was performed in which free phage particles (supernatant after phage adsorption to the host for 12 min) were mixed with 10^8^ CFU/mL E110 and incubated in LB with 2% NaCl. After 4 h of incubation at 37°C, clear zones were recorded.

### Capsule extraction and quantification

The capsule on the surface of *V. alginolyticus* was extracted, and uronic acids were quantified to assess capsule production. *V. alginolyticus* cultures of 50 mL were cultivated in LB broth containing 2% NaCl at 37°C. The cells were collected through centrifugation at 8000 g for 5 min at 4°C and subsequently rinsed using 10 mL of sterile saline solution (2% NaCl). Afterward, the cells were pelleted again with the same centrifugation parameters and were resuspended in 10 mL of 50 mM EDTA (pH 8.0). The sample was incubated at 30°C for 1 h, and the residue was washed with 10 mL of sterile saline solution (2% NaCl). The washed precipitate was gathered through centrifugation at 8000 g for 5 min at 4°C and resuspended in 10 mL of sterile saline solution (2% NaCl), supplemented with 2 mL of 1% 3-sulfopropyltetradecyldimethylbetaine (pH 2.0). After thorough mixing, the sample was heated at 50°C for 30 minutes and centrifuged at 8000 g, 5 min, and 4°C. The supernatant (12 mL) was mixed with a chloroform-n-butanol mixture (5:1, 24 mL) through inversion and at room temperature for 30 min. The mixture was centrifuged at 8000 g, 5 min, and 4°C. The topmost aqueous phase, which contained CPS, was precipitated with four times the cold 100% ethanol volume through inversion and incubated at 4°C for 12 h. The samples were pelleted through centrifugation at 12,000 g for 10 min at 4°C, washed with 70% ethanol, and lyophilized using the vacuum freeze dryer (Alpha-2LDPLUS, Christ).

To quantify purified CPS, 100 μL of the CPS solution was added to 5 μL of pre-cooled 4 M ammonium sulfamate and 600 μL of 25 mM sodium tetraborate. The mixture was vigorously vortexed and boiled for 15 min. After cooling to room temperature, 40 μL of 0.15% 3-hydroxyldiphenol was added, and the samples were divided into aliquots in a 96-well plate. The absorbance was measured at 520 nm, and standard curves were generated using 100 μL of glucuronic acid within a concentration range of 0 to 40 μg/mL.

### AI-2 activity assays

The AI-2 activity bioassay was performed according to the method previously described (47). The cell-free culture fluids of *V. alginolyticus* were prepared using the method described above at 0, 3, 6, 9, 12, and 24 h incubations. *V. harveyi* BB170 was incubated overnight and diluted 1:5000 in fresh AB medium. Add 20 µL of the cell-free culture fluids and 180 µL of the *V. harveyi* BB170 dilution to a 96-well black flat-bottom microtiter plate and incubate at 30°C with shaking at 175 rpm. At the same time, add sterile LB liquid medium in the same proportion to dilute *V. harveyi* BB170 as a negative control, and repeat three times for each sample. Samples were measured and detected at 460 nm every hour until 8 h. The ratio of maximal bioluminescence (test strain)/bioluminescence (negative control) represents the activity of AI-2 produced by the test strain.

### RT-qPCR

Cells were harvested at the indicated OD_600_. RNA was extracted with TRIzol and purified using an RNA Miniprep kit (Qiagen, Germany). Total cDNA was synthesized using the HiScript II reverse transcriptase kit (Vazyme, China). Quantitative real-time reverse transcription PCR (RT-qPCR) was performed using the SYBR Green Real-Time PCR Master Mix and the Step One Plus (ABI, USA) Real-Time PCR System. The 2^−ΔΔCT^ method was used to calculate the relative expression levels of genes using *dnaK* as the reference gene.

### Statistical analysis

All measurement data were expressed as mean ± standard deviation, and differences between groups were evaluated using Student’s t-test for individual measurements or two-way analysis of variance (ANOVA) for data containing repeated measurements of the same cultures. Statistical analysis was done using GraphPad Prism, version 5, software (GraphPad Software, Inc., CA).

## ACKNOWLEDGMENTS

This work was partially supported by the National Natural Science Foundation of China (31872597), Jiangsu Agricultural Science and Technology Independent Innovation Fund (CX [23]1007), Fundamental Research Funds for the Central Universities (B220203038), Postgraduate Research & Practice Innovation Program of Jiangsu Province (42200333).

We thank Mao Lin for kindly providing *V. alginolyticus* VP505, VP506, and *Autographiviridae* phage φVP506and φVP506.

## AUTHOR CONTRIBUTIONS

Xixi Li: Conceptualization, Methodology, Validation, Formal analysis, Writing-original draft; Chen Zhang, Shenao Li, Sixuan Liang, and Xuefei XU: Methodology, Formal analysis; Zhe Zhao: Conceptualization, Formal analysis, Writing-original draft, Writing-review & editing, Resources, Supervision, Funding acquisition. All authors approved the version to be published.

## SUPPLEMENTAL MATERIAL

### Figure Legends

**Figure S1. *V. alginolyticus* E110 quorum-sensing (QS) mutants influence phage infection capacity. (A)** Optical densities (OD_600_) of cultures of E110 wild-type (WT) and QS mutants Δ*luxS* and complemented strain Δ*luxS: luxS* in the presence or absence of phage HH109 at a multiplicity of infection (MOI) of 0.001 were measured in a 96-well microtiter plate containing 200 μL of each culture using the Bioscreen C at every 40-min interval for 520 min. Data are averages of three samples with standard deviations (error bars). **(B)** OD_600_ of cultures of E110 (WT) and QS mutants Δ*luxM* and complemented strain Δ*luxM: luxM* in the presence or absence of phage HH109 at an MOI of 0.001 were measured in a 96-well microtiter plate containing 200 μL of each culture using the Bioscreen C at every 40-min interval for 520 min. Data are averages of three samples with standard deviations (error bars). **(C)** OD_600_ of cultures of E110 (WT) and QS mutants Δ*cqsA*, and complemented strain Δ*cqsA: cqsA* in the presence or absence of phage HH109 at an MOI of 0.001 was measured in a 96-well microtiter plate containing 200 μL of each culture using the Bioscreen C at every 40-min interval for 520 min. Data are averages of three samples with standard deviations (error bars). **(D)** OD_600_ of cultures of E110 (WT) and QS mutants Δ*hnoX* and complemented strain Δ*hnoX: hnoX* in the presence or absence of phage HH109 at an MOI of 0.001 were measured in a 96-well microtiter plate containing 200 μL of each culture using the Bioscreen C at every 40-min interval for 520 min. Data are averages of three samples with standard deviations (error bars).

**Figure S2. Effect of the quorum sensing regulator AphA on *Autographiviridae* phage infection in *V. alginolyticus* strains. (A)** Optical densities (OD_600_) of cultures of E110 wild-type (WT) and mutants Δ*aphA* and complemented strain Δ*aphA:*Δ*aphA* in the presence or absence of phage HH109 at a multiplicity of infection (MOI) of 0.001 were measured in a 96-well microtiter plate containing 200 μL of each culture using the Bioscreen C at every 20-min interval for 520 min. Data are averages of three samples with standard deviations (error bars). **(B)** OD_600_ of cultures of VP505 and QS mutants VP505Δ*aphA* and complemented strain VP505Δ*aphA*:Δ*aphA* in the presence or absence of phage φVP505 at a multiplicity of infection (MOI) of 0.01 was measured in a 96-well microtiter plate containing 200 μL of each culture using the Bioscreen C at every 20-min interval for 12 h. Data are averages of three samples with standard deviations (error bars). **(C)** OD_600_ of cultures of VP506 and QS mutants VP506Δ*aphA* and complemented strain VP505Δ*aphA: aphA* in the presence or absence of phage φVP506 at a multiplicity of infection (MOI) of 0.001 was measured in a 96-well microtiter plate containing 200 μL of each culture using the Bioscreen C at every 20-min interval for 12 h. Data are averages of three samples with standard deviations (error bars). **(D)** Adsorption rate of phage HH109 by its host strains *V. alginolyticus* E110 (WT) and mutants Δ*aphA* and complemented strain Δ*aphA: aphA* at every 6-min interval for 12 h. Data are averages of three samples with standard deviations (error bars). **(E)** Adsorption rate of phage φVP505 by its host strains *V. alginolyticus* VP505 and QS mutants VP505Δ*aphA* and complemented strain VP505Δ*aphA: aphA* at every 3-min interval for 6 min. Data are averages of three samples with standard deviations (error bars). **(F)** Adsorption rate of phage φVP506 by its host strains *V. alginolyticus* VP506 and QS mutants VP506Δ*aphA* and complemented strain VP506Δ*aphA: aphA* at every 3-min interval for 9 min. Data are averages of three samples with standard deviations (error bars).

**Figure S3. QS activates *ugd* expression. (A)** The expression of critical genes of CPS at high cell density in VP506 or VP505 and the designated QS mutants. **(B)** Relative *ugd* expression was measured by RT-qPCR in *V. alginolyticus* VP506 or VP505 cells at low and high cell densities (OD_600_, 0.8, and 2.5, respectively). The reference gene was *dnaK*. Data are averages of three samples with standard deviations (error bars). ***, *P* < 0.01 (paired *t* test).

**Figure S4. AI-2 signal molecular production *o*f *V. alginolyticus* in different growth phases. (A)** AI-2 activity of *V. alginolyticus* E110 at 30℃. Levels of AI-2 production were shown in logarithmic form. **(B)** AI-2 activity of *V. alginolyticus* VP505 at 30℃. Levels of AI-2 production were conducted in logarithmic form. **(C)** AI-2 activity of *V. alginolyticus* VP506 at 30℃. Levels of AI-2 production were shown in logarithmic form.

### Table Legends

**Table S1. Bacterial strains and plasmids used in this study.**

**Table S2. Primers used in this study.**

